# Machine Learning-Driven fMRI Analysis for Objective Craving Prediction

**DOI:** 10.1101/2025.04.17.649130

**Authors:** Hajar Mahdavi-Doost, Ghazaleh Soleimani, Kelvin O Lim, Hamed Ekhtiari

## Abstract

**Background:** Craving is a fundamental aspect of substance use disorder (SUD), traditionally assessed through subjective self-report measures. To develop more objective assessments, we created a brain-based marker to predict craving based on machine learning approaches using functional magnetic resonance imaging (fMRI) drug cue reactivity data from 69 participants with methamphetamine use disorders.

**Methods:** To predict craving intensity, our analysis demonstrated that utilizing principal component analysis (PCA) and linear regression outperformed other models in terms of Root Mean Squared Error (RMSE). Employing a 5-fold cross-validation strategy with a 20% holdout set, we established the reliability of the model. Additionally, the model successfully classified high and low craving levels and distinguished cue types (neutral vs. drug) based on fMRI data.

**Results:** The model achieved an RMSE of 0.983 ± 0.026 (standard deviation), with strong generalization evidenced by an out-of-sample RMSE of 0.985 and statistical significance (*p* < 0.026; effect size (Cliff’s Delta) = 0.715; statistical power = 0.639). Key neurobiological signatures included the Parahippocampal Gyrus, Superior Temporal Gyrus, Medioventral Occipital Cortex, and Amygdala (positively associated with craving), as well as the Inferior Temporal Gyrus (negatively associated). Classification of high vs. low craving levels yielded an AUC-ROC of 0.684 ± 0.084 (out-of-sample AUC-ROC = 0.714), with significant separation (*p* < 0.04; Cliff’s Delta = 0.831). Additionally, classification of cue types (neutral vs. drug) achieved an AUC-ROC of 0.692 ± 0.090 (out-of-sample AUC-ROC = 0.693), with *p* < 0.002, Cliff’s Delta = 0.896, and statistical power = 0.800, highlighting the robustness of the model.

**Conclusion:** These findings underscore the potential of neuroimaging and machine learning to provide objective, data-driven insights into the neural mechanisms underlying subjective experience of craving and to inform future clinical applications in SUD.

## 1. Introduction

Craving, characterized as an intense desire to consume drugs or food, is a fundamental driver of substance use disorder (SUD), contributing to the compulsive drive to seek drugs despite negative consequences (Koob & Volkow, 2010; Tiffany & Wray, 2012). It is a potent predictor of relapse (Sinha, 2013; Weiss et al., 1995) and has thus become a major focus of addiction research. Functional MRI of drug cue reactivity (FDCR) is a technique in addiction research to understand how the brain responds to drug-related cues. It involves exposing individuals to drug-related stimuli (such as images, sounds, or even drug) while capturing brain activity using fMRI. The goal is to identify neural circuits that are activated in response to these cues, as they are associated with craving and relapse in substance use disorders (Addiction Cue-Reactivity Initiative (ACRI) Network et al., 2024; Ekhtiari et al., 2022; Pollard et al., 2023). This method has also been expanded to explore the brain’s response to food-related cues, demonstrating its utility in studying craving mechanisms across substance and non-substance domains (Field & Cox, 2008; Koban et al., 2023).

Research in neuroscience has consistently shown that drug cues elicit activity in several key brain regions. They include the ventromedial prefrontal cortex (vmPFC) and ventral striatum which are known for their roles in reward processing and decision-making, and the amygdala and hippocampus which are crucial for emotional regulation and memory, especially in response to drug cues (Kilts et al., 2004; Koob & Volkow, 2010; Volkow et al., 1997; Zilverstand et al., 2018). The insula has been known for its involvement in interoceptive awareness, helping individuals detect internal bodily states associated with craving (Naqvi & Bechara, 2010). These regions collectively form the foundation of the neurocircuitry of addiction, emphasizing both the sensory and emotional components of craving (Koob & Volkow, 2010; Naqvi & Bechara, 2010; Volkow et al., 2016).

Despite these advancements, our understanding of the neural basis of craving remains incomplete. Current neuroimaging literature highlights the involvement of various brain regions in craving (Volkow et al., 2016; Zilverstand et al., 2018; Seo et al., 2011). However, many studies fail to implement robust predictive modeling techniques, such as cross-validation, which limits the generalizability of their findings. Instead, they often rely on correlation analyses between subjective craving scores and activity or connectivity in specific brain regions, which are prone to overfitting and circularity in feature selection (Kriegeskorte et al., 2009). These limitations undermine the reliability of population-level inferences and hinder the development of practical neuroimaging-based biomarkers. Machine learning approaches offer a solution by enabling the integration of high-dimensional brain activity patterns into predictive models that can generalize across individuals and contexts. Proper cross-validation within these models is essential for testing predictive power, increasing replicability, and facilitating the translation of neuroimaging findings into real-world applications, such as diagnostic tools or personalized treatment interventions. This underscores the urgent need for data-driven methodologies that leverage machine learning to provide sensitive, specific, and generalizable measures of craving.

Recently, there has been significant progress in leveraging machine learning techniques for craving prediction. Some studies utilized connectome-based predictive modeling (CPM), a machine learning framework designed to identify functional connectivity patterns predictive of craving in various addictions (Antons et al., 2023; Garrison et al., 2023; Shen et al., 2017). Unlike traditional correlation-based analyses, CPM employs robust cross-validation to ensure the generalizability of its findings, making it a more reliable approach for understanding the network-level mechanisms of craving. Similarly, Koban et al. (2023) employed LASSO-PCR (Least Absolute Shrinkage and Selection Operator – Principal Component Regression) to develop the Neurobiological Craving Signature (NCS). This activity-based approach identifies a reproducible pattern of brain activity involving the ventromedial prefrontal cortex, ventral striatum, temporal/parietal association areas, cingulate cortex, and mediodorsal thalamus. Their study covered individuals with cocaine use disorder, alcohol use disorder, and smoking addiction, using drug and food-related visual cues to train and validate their models. The use of study-stratified ten-fold cross-validation ensured robust predictive accuracy, addressing some limitations of traditional methods. While these studies highlight the potential of machine learning to uncover reproducible brain-based markers of craving, their generalizability across specific substances, such as methamphetamine, and individual variability remains an area for further development.

Our current study seeks to build upon this body of work by using fMRI drug cue reactivity from 69 participants with methamphetamine use disorder (MUD) and employing machine learning models to train a predictor being able to predict subjective craving levels based objective neuroimaging data. This will enable us to create detailed maps of brain activity that predicts subjective response to drug cues. These findings could provide opportunities to use fMRI markers as valid measure of craving to monitor response to various interventions and inform targeted interventions for SUD such as transcranial magnetic stimulation (TMS)(R. Chen et al., 1997; Hanlon et al., 2015; Jansen et al., 2015; Pettorruso et al., 2020), transcranial direct current stimulation (tDCS) (Chan et al., 2024; J. Chen et al., 2020) and deep brain stimulation (DBS) (Kuhn et al., 2014; Luigjes et al., 2012).

## 2. Methods

This section outlines the procedures for data collection, imaging protocols, and the analytical framework applied to predict craving intensity from fMRI data. Details on participant selection, imaging settings, data preprocessing, and statistical analyses are provided in the following sections.

### 2.1 Participants

A total of 69 male participants (mean age ± one standard deviation = 35.86 ± 8.47 years), all diagnosed with methamphetamine use disorder in their abstinence phase, were recruited for this study. These participants were admitted to the 12&12 Center, a residential abstinence-based treatment facility in Tulsa, Oklahoma. They were either part of the residential program or its aftercare transitional living programs. They underwent the fMRI drug cue reactivity task at Laureate Institute for Brain Research (LIBR), in Tulsa, OK.

The study included participants who met the following conditions: (1) they spoke English, (2) had been diagnosed with methamphetamine use disorder within the last year, (3) were enrolled in a residential program designed to support abstinence from methamphetamine, (4) had maintained at least one week of abstinence prior to the study, and (5) demonstrated the ability and willingness to engage in the informed consent process.

Individuals were excluded if they (1) were unwilling or unable to complete essential parts of the study, including MRI scanning (e.g., due to claustrophobia), drug cue rating, or behavioral evaluations; (2) reported over six months of methamphetamine abstinence; (3) had a diagnosis of schizophrenia or bipolar disorder based on a MINI interview; (4) exhibited active suicidal thoughts, determined either by self-report or through assessment by study staff at any stage of the study; or (5) had a positive screening for amphetamines, opioids, cannabis, alcohol, phencyclidine, or cocaine, as confirmed by breath or urine testing.

### 2.2 fMRI Task

All participants in this study completed the same fMRI drug cue reactivity task structure as described in (Ekhtiari et al., 2016). During the fMRI task, participants were shown alternating blocks of drug-related (methamphetamine) and neutral cues. Each block consisted of six images, each displayed for five seconds with a 200ms blank screen between them, making the total duration of each block 31 seconds. After each block, participants rated their craving based on the question “How much craving do you have right now?”. The craving scale was from one to four, with one indicating “No Urge” and four indicating “Strong Urge.” The interval between blocks varied between 8 and 12 seconds, and the blocks alternated between drug and neutral, starting with a neutral block. A total of eight blocks were presented, with four of each type. The total scan time for this task was approximately 6.5 minutes (Figure 1).

**Figure 1.**
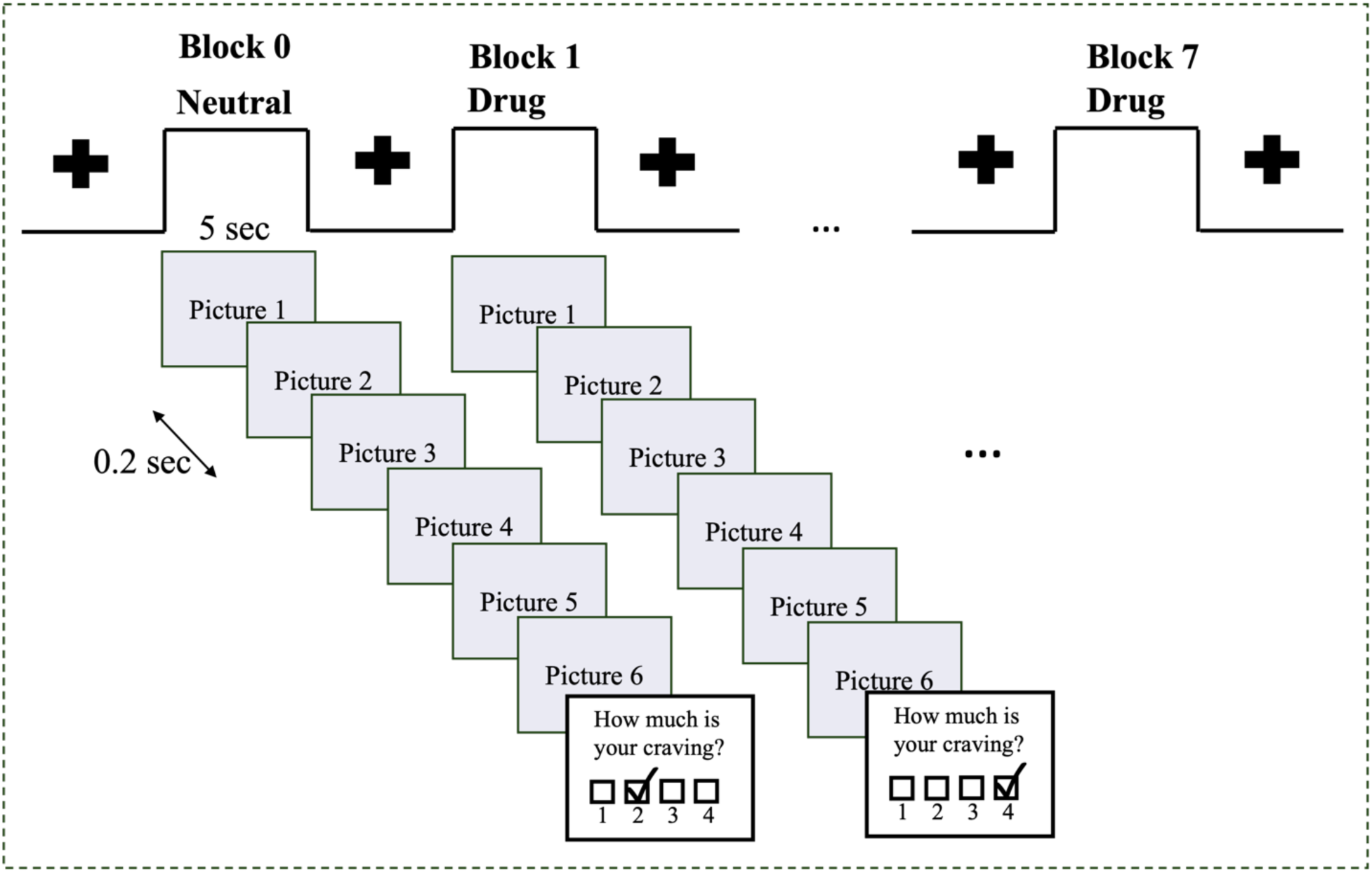
fMRI drug cue reactivity task. The task included alternative blocks of neutral and drug cues (images). Each block consisted of 6 images each displayed for 5 seconds with a 0.2 sec inter-stimulus interval resulting in a total block duration of 31 seconds. A visual fixation point was presented for 8 to 12 sec between consecutive blocks. At the end of each block a self-report was collected which participants were asked to rate their current meth craving level on a 1 to 4 rating scale (1 lowest to 4 highest).

### 2.3 Image Acquisition

The MR images were collected using two General Electric (GE) MRI 750 3T scanners located at LIBR. The FDCR task lasted 6 minutes and 32 seconds, with scan parameters as follows: Repetition Time/Echo Time (TR/TE) = 2000/27ms, Field of View (FOV)/slice=240/2.9mm,128× 128 matrix resulting in 1.875 × 1.875 × 2.9 mm voxels, across 39 axial slices with 196 repetitions. High-resolution structural images were obtained using an axial T1-weighted Magnetization Prepared Rapid Gradient Echo (MPRAGE) sequence, with parameters including TR/TE=5/2.012ms, FOV/slice=240× 192/0.9mm, 256× 256 matrix generating 0.938 × 0.928 × 0.9 mm voxels, and 186 axial slices.

### 2.4 fMRI Data Preprocessing

First-level processing was performed using Analysis of Functional NeuroImages (AFNI), which involved several steps: discarding the first three pre-steady state images, despiking, correcting for slice timing, realigning the images, transforming them to MNI space, and applying 4 mm Gaussian full-width-half-maximum (FWHM) smoothing. Following this, regression analysis was conducted, incorporating nuisance regressors for the first three polynomial terms and the six motion parameters. TRs with the Euclidean norm of the derivative of the six motion parameters being greater than 0.3 points were removed. After preprocessing, a General Linear Model (GLM) was applied to obtain the beta coefficients. This processed data was subsequently used to train a machine learning algorithm.

### 2.5 Data Characteristics

The beta coefficients derived from the fMRI data presented several analytical challenges. These coefficients, which estimate neural activity associated with specific task conditions, exhibited no readily discernible patterns, making it difficult to identify meaningful clusters or activation profiles (see Supplementary Figure 1). Additionally, the dataset was characterized by a relatively small sample size and extremely high dimensionality (over one million voxels), with a substantial proportion of zero values—further complicating the analysis. Craving ratings showed clear differentiation between drug and neutral cues, with overall moderate variability. Detailed summary statistics are presented in Supplementary Figure 2.

### 2.6. Data Analysis Pipeline

In our data analysis pipeline (Figure 2), we employed several robust methods to ensure the reliability and validity of our results. We utilized 5-fold cross-validation while making sure that the splitting of data to train and test was done in a subject-level manner. In other words, we made sure the data belonging to the same subject was assigned to the same split. This avoids leaking the subjects’ information to other splits and makes sure a reliable model development (Kohavi, 1995). Additionally, we used hold-out testing by setting aside 20 percent of the data as a hold-out test set to evaluate the model’s performance on unseen data.

**Figure 2.**
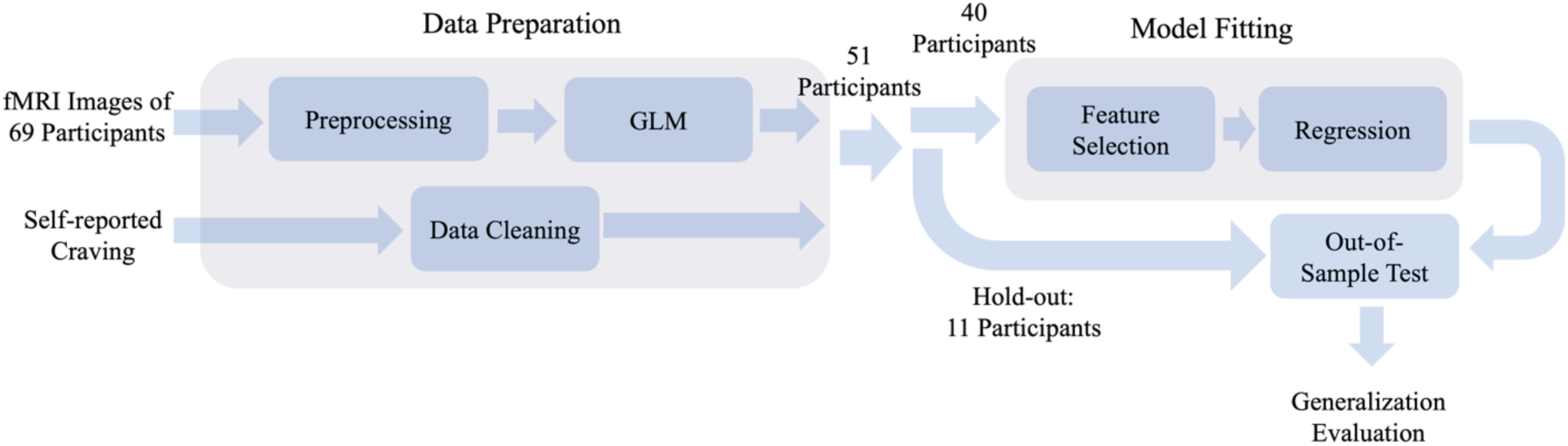
Analysis pipeline. fMRI images and self-reported craving levels from 69 participants are processed in the data preparation unit. The fMRI data is preprocessed, and a general linear model is applied to generate beta coefficients for data analysis. At the same time, participants with missing data (17 participants) are removed. Subsequently, 20% of the data (11 participants) is held out for testing, while the model is fitted using the remaining 80%, comprising approximately 40 participants. The model fitting includes a feature selection step, where the high-dimensional data is projected to a lower dimension, followed by training a regression model. Finally, the model is tested on the unseen portion of data to evaluate its generalizability.

To assess the model performance across varying degrees of craving intensity, we stratified metrics based on different craving levels. In other words, the performance was evaluated separately for each craving level and then the results were averaged to obtain an overall performance score. This way, we don’t have to worry about the craving levels being imbalanced in each split as we already used the stratified metrics. We initially explored a mixed-effects model to account for within-subject variability across multiple trials per participant. However, the objective of machine learning—where models are trained to generalize to unseen subjects— fundamentally differs from the intent of mixed-effects modeling, which explicitly captures subject-specific variations. The limited per-subject sample size and minimal variance explained by random effects further reduced its effectiveness. In contrast, standard machine learning approaches, such as linear regression, optimize fixed effects across the entire dataset, enhancing generalizability and predictive performance for new subjects.

We also conducted permutation test to evaluate the significance of our results, involving the repeated shuffling of craving levels (1000 times) to generate a distribution of outcomes under the null hypothesis, which allowed us to determine the statistical significance of our findings. These approaches were crucial in developing a reliable and generalizable predictive model for craving intensity.

### 2.7 The Model and Hyperparameter Tuning

We have implemented a comprehensive two-step process for feature selection, followed by model training (Figure 2). Initially, we evaluated PCA and Analysis of Variance (ANOVA) for feature selection. The features correspond to distinct voxel locations within the brain. Subsequently, we assessed 5 regression algorithms, including Multivariate Linear Regression (Anderson, 1958), Ridge Regression (Hoerl & Kennard, 1970), Lasso Regression (Tibshirani, 1996), Elastic Net (Zou & Hastie, 2005), Random Forest (Breiman, 2001), and XGBoost (T. Chen & Guestrin, 2016), to identify the optimal model that minimizes RMSE. These methods were selected based on their complementary strengths in handling high-dimensional data and regularization. Multivariate Linear Regression serves as a baseline model to evaluate the effectiveness of other approaches. Ridge and Lasso Regression are well-suited for reducing overfitting in high-dimensional data through L2 and L1 regularization, respectively. Elastic Net combines these two penalties to handle correlated features effectively. Random Forest, a robust ensemble learning method, was chosen for its ability to model non-linear relationships and inherent feature importance evaluation. Finally, XGBoost, a state-of-the-art gradient boosting algorithm, was included for its efficiency and strong performance in a wide range of regression tasks. Moreover, before feature selection, we eliminate features with very low variance (Var < 0.001). Additionally, we ensure that the data is centered using a standard scaler before applying PCA. To determine the optimal hyperparameters, we employ the GridSearchCV (Pedregosa et al., 2011) class from the scikit-learn Python library, calculating RMSE for different scenarios (see Supplementary Table 1).

In the optimized pipeline, the data used in model fitting was projected into a lower-dimensional space by multiplying it with the eigenvector matrix (V) derived from PCA. This projected data was then combined with the linear regression coefficient vector (B) to predict craving levels. The resulting product VB represents a key neurobiological signature of craving (Koban et al., 2023) forming a one-dimensional vector that indicates the contribution of each voxel to the craving process. To identify the activated subregions more closely, Brainnetome atlas parcellation was applied to extract 246 subregions, each containing a cluster of voxels (Fan et al., 2016). Each voxel was assigned to its corresponding Brainnetome subregion, and the average VB value for all voxels within each subregion was calculated.

In general, the craving intensity, particularly whether it is high or low is of primary interest. To enhance model interpretability, we classified craving levels into high and low and focused on the most extreme craving states. Here, the steps of the algorithm align with those implemented in the regression algorithm, with an additional mapping step. Here, the proper number of PCA components was determined to be 50 due to lower number of samples. During the mapping phase, the regression algorithm’s outcomes are categorized into high or low levels, reflecting the predicted outcomes. Given that the predicted results of the regression are not uniformly distributed between values of 1 and 4, and that this distribution varies in each iteration of cross validation, we need to optimize the threshold level for the classification. We utilized F_1_ score as the metric for threshold selection due to the label imbalance observed in each iteration, ultimately selecting the threshold that maximizes the F_1_ score at each iteration.

To evaluate the robustness of our algorithm, we trained a predictor to classify cue types as neutral or drug-related (defined as 0 and 1, respectively). We again employed the same pipeline as before with the addition of a step mapping the regression output to 0 and 1 with the threshold equal to 0.5 and the number of components of PCA equal to 100.

## 3. Results

We have considered 3 primary aims for this project. First, aimed to predict the intensity of craving. Following the pipeline in Figure 2, we optimized the model and proved that the result is statistically significant. This identified a brain map highlighting the contribution portion of each subregion. Next, we developed a classification algorithm to be able to distinguish between the high and low craving based on fMRI images and proved its statistical significance. Lastly, we developed an algorithm differentiating the cue types and again confirmed its significance.

### 3.1 Craving Intensity Prediction

Our analysis shows that linear models, including Linear Regression, Lasso, and Elastic Net, with slightly similar performance, outperform non-linear models in both feature selection (ANOVA) and regression algorithms (Random Forest and XGBoost). In conclusion, PCA with 100 components and linear regression emerged as the best approach for this analysis (Supplementary Table 1).

The results presented in Table 1 demonstrate the statistical comparison between the Proposed Algorithm and the Null Hypothesis across three key performance metrics: RMSE, MAE, and Correlation Coefficient. The Shapiro-Wilk test was used to assess normality, revealing that the correlation coefficient followed a normal distribution, while RMSE and MAE did not. Given this, Cohen’s d was used as the effect size measure for correlation, and Student’s t-test was employed to compute the confidence interval (95% CI = [0.209, 0.222]). The corresponding p-value (0.050) and effect size (1.826) indicate a strong difference between the two methods, with a statistical power of 50.5%.

Conversely, RMSE and MAE exhibited non-normal distributions, necessitating the use of Cliff’s Delta for effect size and bootstrap resampling for confidence interval estimation. For RMSE, the 95% CI [0.980, 0.984] was obtained via bootstrapping, with an effect size of 0.715, a p-value of 0.028, and a statistical power of 63.9%. Similarly, for MAE, the 95% CI [0.939, 0.941] was computed through bootstrapping, with an effect size of 0.836, a p-value of 0.008, and a statistical power of 74.6%.

Additionally, statistical significance (p-value) and power were determined by counting from the 5% critical value threshold to evaluate the reliability of the observed differences. The out-of-sample performance further supports these findings, with RMSE and MAE showing robust generalization (0.985 and 0.976, respectively), whereas correlation exhibited a lower out-of-sample predictive value (0.108).

Supplementary Figure 3 outlines the out-of-sample results for the regression problem. For each participant, only the minimum and the maximum self-reported craving levels were considered. In cases with multiple instances of minimum or maximum cravings, only the first occurrence was used. The fitted normal distribution of the outcomes and associated box plots are presented in Figure 3.

The neurobiological signature of craving associated to our algorithm is illustrated in Figure 4. Regions with higher values amplify the corresponding beta coefficients, while regions with lower values attenuate them. For visualization purposes, the signature was normalized to align with the norm of the average beta coefficients derived from the fMRI data. Figure 5 displays the Brainnetome bar plot, showing that the Parahippocampal Gyrus, Superior Temporal Gyrus, Medioventral Occipital Cortex, and Amygdala contributed most to craving, while the Inferior Temporal Gyrus exhibited the most negative contribution.

**Table 1.** Statistical comparison of the proposed algorithm and the null hypothesis for the prediction of craving level. The mean, standard deviation (Std Dev), 95% confidence interval (CI), p-value, effect size, and statistical power are reported. The out-of-sample performance is also included to evaluate generalization. Lower RMSE and MAE values indicate better predictive performance, while higher Pearson correlation show better performance.

| Metric | Mean | Std Dev | 95% CI | P-value | Effect Size | Statistical Power (%) | Out-of-Sample |
| --- | --- | --- | --- | --- | --- | --- | --- |
| RMSE | 0.983 | 0.026 | [0.98 , 0.984] | 0.028 | 0.715 | 63.9 | 0.985 |
| MAE | 0.94 | 0.014 | [0.939 , 0.941] | 0.008 | 0.836 | 74.6 | 0.976 |
| Correlation | 0.216 | 0.112 | [0.209 , 0.222] | 0.05 | 1.826 | 50.5 | 0.108 |

**Figure 3.**
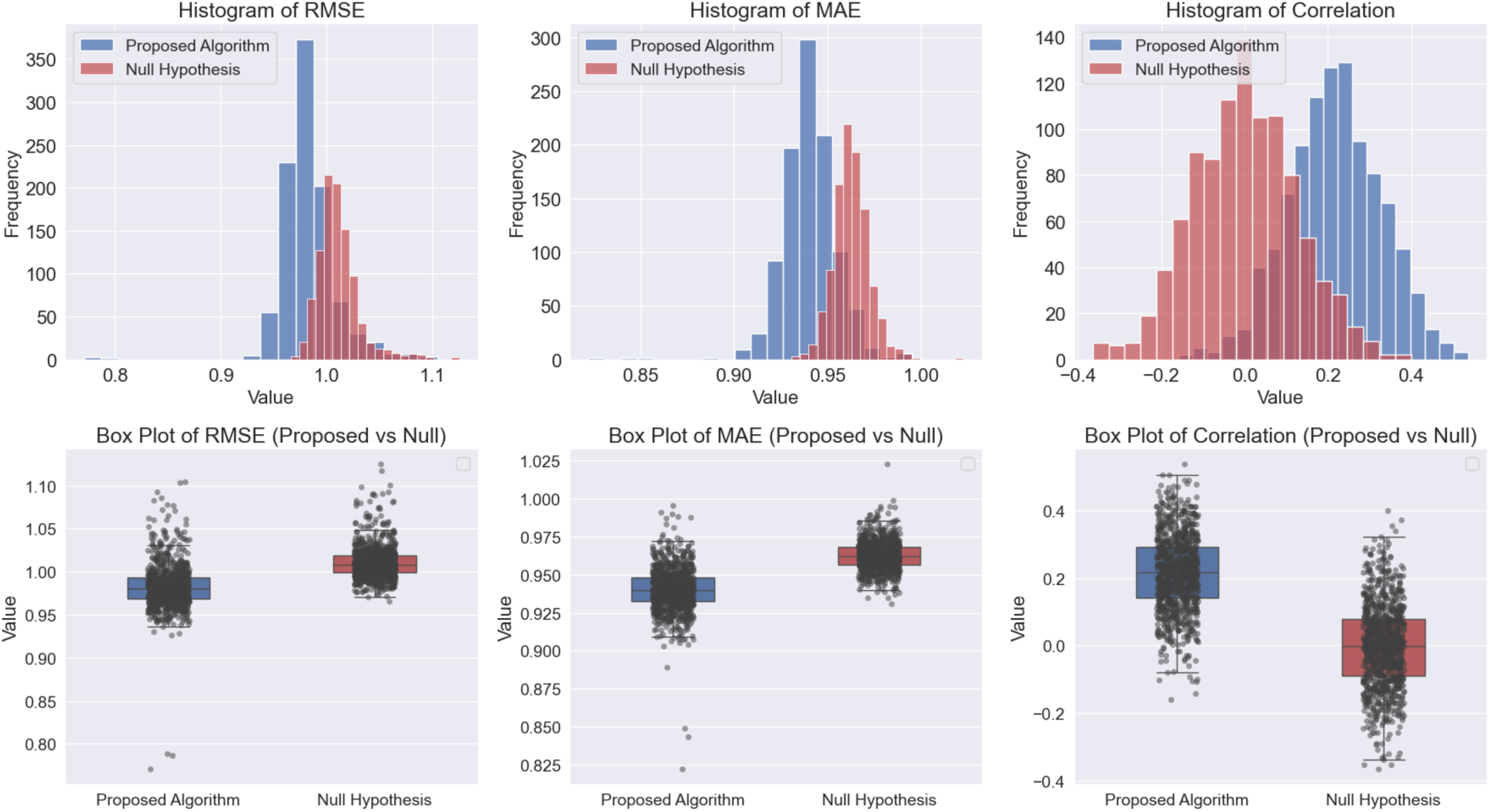
Comparison of the results between the proposed algorithm and the null hypothesis across three evaluation metrics: RMSE, MAE, and correlation coefficient. Top Row: The histograms illustrate the distributions of results for the proposed algorithm (blue) and the null hypothesis (red). Bottom Row: The box plots provide a comparative visualization of distribution spread and variability. Each box represents the interquartile range (IQR) (middle 50% of data), with the central line indicating the median. The whiskers extend up to 1.5 times the IQR, and data points beyond these whiskers are considered outliers, plotted as individual dots.

**Figure 4.**
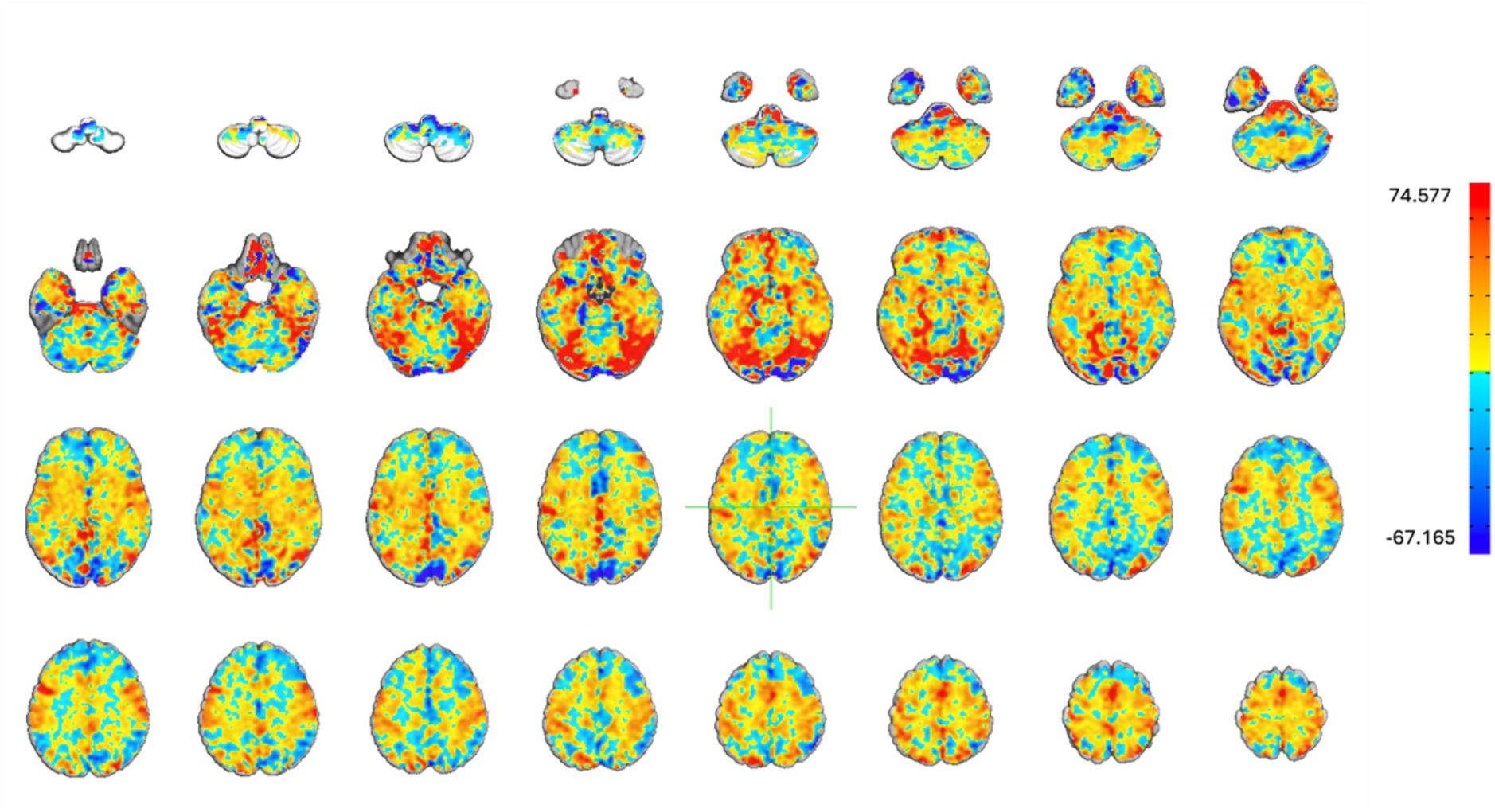
Neurobiological signature of craving. This map illustrates the contribution of each region to the prediction of subjective craving intensity. The red regions represent areas with positive contributions, while the blue regions indicate areas with negative contributions.

**Figure 5.**
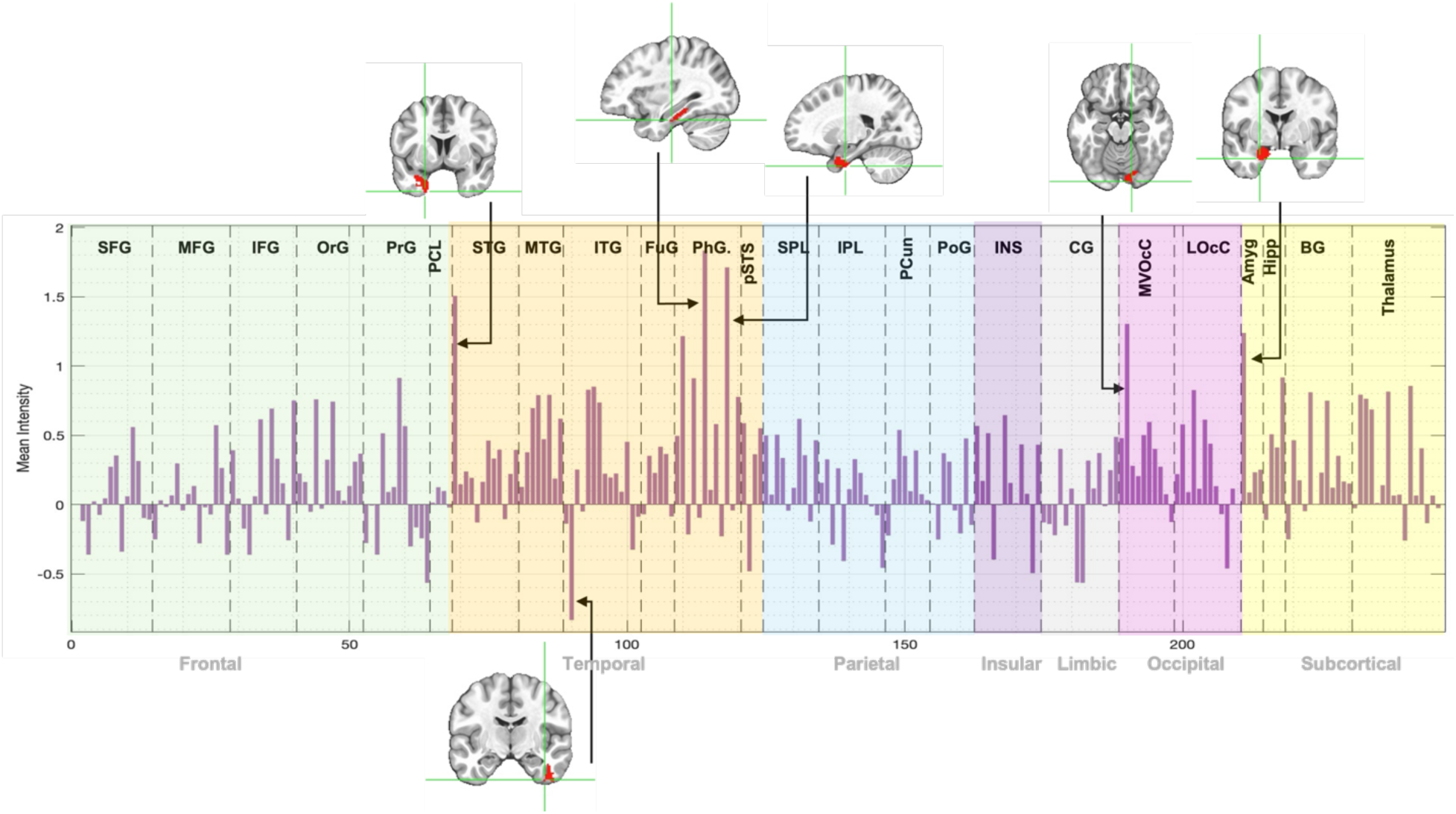
Brainnetome Subregions Bar Plot. An illustration of the involvement of various brain subregions from the Brainnetome Atlas in relation to craving. Each bar represents a specific subregion, with the height indicating the strength of contribution with craving. Key neurobiological signatures identified were the Parahippocampal Gyrus, Superior Temporal Gyrus, Medioventral Occipital Cortex, and Amygdala, which showed positive contributions to craving, and the Inferior Temporal Gyrus, which exhibited a negative contribution to craving.

### 3.2 High and Low Craving Classification

The performance of the proposed algorithm compared to the null hypothesis is presented in Table 2. RMSE, accuracy, and area under the receiver operating characteristic curve (AUC-ROC) were evaluated to assess prediction and classification performance.

For classification accuracy, the proposed algorithm demonstrates some improvements in the evaluated metrics compared to the null hypothesis. It is important to note that all metric distributions follow a non-normal distribution; therefore, we employ non-parametric or appropriate statistical methods for estimating parameters. The accuracy of 0.674 ± 0.108 indicates that the algorithm’s performance exceeds chance level with a moderate margin. The confidence interval of [0.668, 0.681], calculated using bootstrap sampling, supports the reliability of the results though the p-value of 0.07 suggests moderate statistical significance. The effect size, measured by Cliff’s Delta (0.753), indicates a substantial difference; however, the limited statistical power (30.3%) underscores the need for further validation with larger sample sizes. The out-of-sample accuracy was 0.703, surpassing the in-sample accuracy, which suggests strong generalization to unseen data.

The AUC-ROC, which evaluates the classifier’s ability to distinguish between low and high craving states, was 0.684± 0.084 for the proposed algorithm. The confidence interval of [0.679, 0.689] confirms the robustness of this metric. The p-value of 0.04 indicates statistical significance, and the effect size (Cliff’s Delta: 0.831) suggests a strong improvement. The statistical power was 43.3%, indicating a limited probability of detecting a true effect with the current sample size. Nonetheless, the out-of-sample AUC of 0.714 provides additional support for the model’s ability to generalize to unseen data.

Overall, the proposed algorithm consistently outperformed the null hypothesis in prediction and classification tasks. The significant improvements in RMSE, accuracy, and AUC-ROC, along with moderate-to-large effect sizes, indicate meaningful differences between the proposed model and the permutation-based null hypothesis. The higher out-of-sample accuracy and AUC further validate the model’s generalization ability in craving classification and prediction.

**Table 2.**
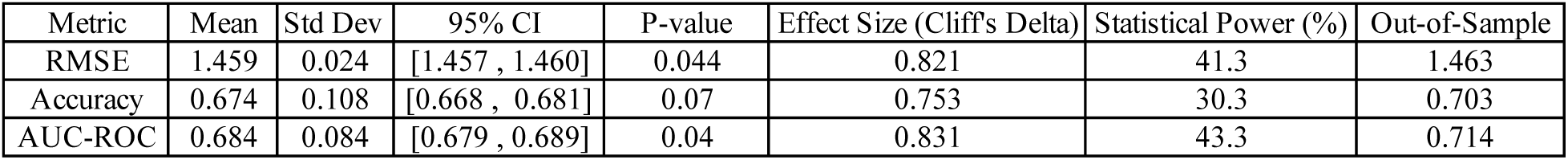
Statistical comparison of the proposed algorithm and the null hypothesis for the classification of high and low craving across RMSE, accuracy, and AUC-ROC metrics. The mean, standard deviation, 95% confidence interval (CI), p-value, Cliff’s Delta effect size, and statistical power are reported. The out-of-sample performance is also included to evaluate generalization. Lower RMSE values indicate better predictive performance, while higher accuracy and AUC-ROC values reflect improved classification capability.

| Metric | Mean | Std Dev | 95% CI | P-value | Effect Size (Cliff's Delta) | Statistical Power (%) | Out-of-Sample |
| --- | --- | --- | --- | --- | --- | --- | --- |
| RMSE | 1.459 | 0.024 | [1.457 , 1.460] | 0.044 | 0.821 | 41.3 | 1.463 |
| Accuracy | 0.674 | 0.108 | [0.668 , 0.681] | 0.07 | 0.753 | 30.3 | 0.703 |
| AUC-ROC | 0.684 | 0.084 | [0.679 , 0.689] | 0.04 | 0.831 | 43.3 | 0.714 |

**Figure 6.**
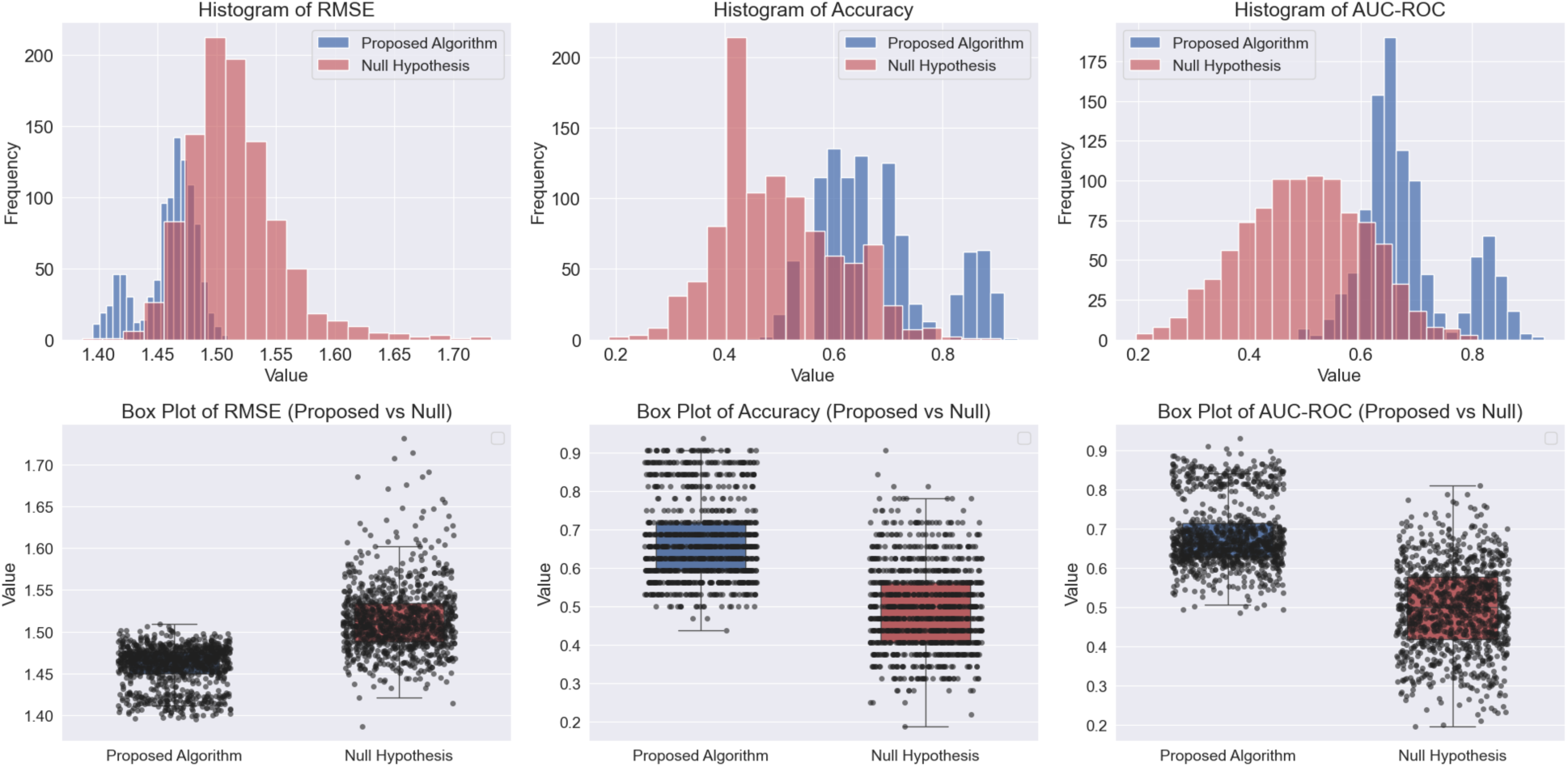
Illustration of the results of classification algorithm for low and high craving levels. Comparison of the proposed algorithm and the null hypothesis (permutation test) for RMSE, accuracy, and AUC-ROC. The top row displays histograms showing the distribution of each metric for both models, with the proposed algorithm in blue and the null hypothesis in red. The bottom row presents box plots illustrating the variability and spread of each metric. The results indicate that the proposed algorithm consistently outperforms the null hypothesis, with lower RMSE and higher accuracy and AUC-ROC values.

The ROC curve (Figure 7) shows the model’s performance on unseen data, with an AUC of 0.70, suggesting moderate discriminatory ability between classes.

**Figure 7.**
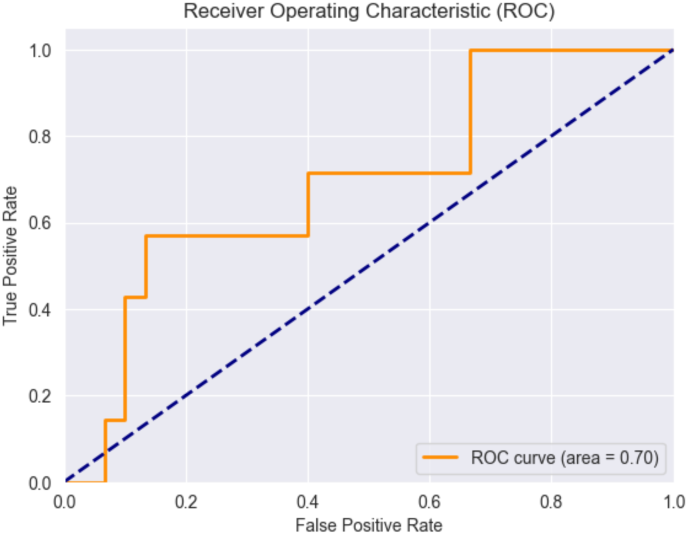
ROC Curve for Classification on Unseen Data. The curve illustrates the trade-off between the True Positive Rate (TPR) and False Positive Rate (FPR) across classification thresholds. The dashed diagonal line represents the baseline performance of a random classifier. The AUC value of 0.70 reflects moderate discriminative ability. The staircase-like shape of the curve is due to the small number of participants in the test set, affecting the curve’s granularity.

### 3.3 Cue Type Classification

Table 3 provides a detailed statistical summary of classifier performance. Since the metrics exhibited a non-normal distribution, we used Cliff’s Delta to estimate effect size and applied bootstrap resampling to compute confidence intervals. The classifier achieves an accuracy of 0.617 ± 0.057, with a Cliff’s Delta of 0.863, indicating a substantial effect size. The highest performance is observed in the precision metric (0.723 ± 0.095, Cliff’s Delta = 0.900), suggesting that when the classifier identifies a cue type, it is often correct. The AUC-ROC score of 0.692 ± 0.090 and AUC-PR of 0.698 ± 0.082 further confirm the model’s strong ability to discriminate between cue types. Notably, the p-values across all metrics are below 0.05, indicating that the observed differences are unlikely due to chance. The statistical power for these tests ranges from 60.5% to 80.0%, reducing the likelihood of Type II errors.

**Table 3.** Cue type classification results. Performance metrics of the cue type classifier, including Accuracy, Precision, AUC-ROC, and AUC-PR. The results demonstrate the classifier’s strong discriminative ability, with significantly higher performance compared to the null hypothesis. The low p-value, high power, and large effect size further support the statistical significance and generalizability of the model.

| Metric | Mean | Std Dev | 95% CI | P-value | Effect Size (Cliff's Delta) | Statistical Power (%) | Out-of-Sample |
| --- | --- | --- | --- | --- | --- | --- | --- |
| Accuracy | 0.617 | 0.057 | [0.613 , 0.620] | 0.02 | 0.863 | 60.5 | 0.614 |
| Precision | 0.723 | 0.095 | [0.717 , 0.729] | 0.02 | 0.9 | 68.1 | 0.727 |
| AUC-ROC | 0.692 | 0.09 | [0.686 , 0.697] | 0.002 | 0.896 | 80 | 0.693 |
| AUC-PR | 0.698 | 0.082 | [0.693 , 0.703] | 0.015 | 0.888 | 64.8 | 0.681 |

The discriminative power and statistical significance of the classifier, assessed using a permutation test against the null hypothesis, are presented in Figure 8 for various performance metrics. The histograms and box plots illustrate the differences between the proposed algorithm and the null hypothesis across accuracy, precision, AUC-ROC, and Area Under the Precision-Recall Curve (AUC-PR). The proposed algorithm consistently outperforms the null hypothesis, as seen in the separation of distributions and higher median values across all metrics. Figure 9 illustrates the ROC curve for the classifier on unseen data, demonstrating a moderate ability to distinguish between cue types, with an AUC-ROC of 0.69.

**Figure 8.**
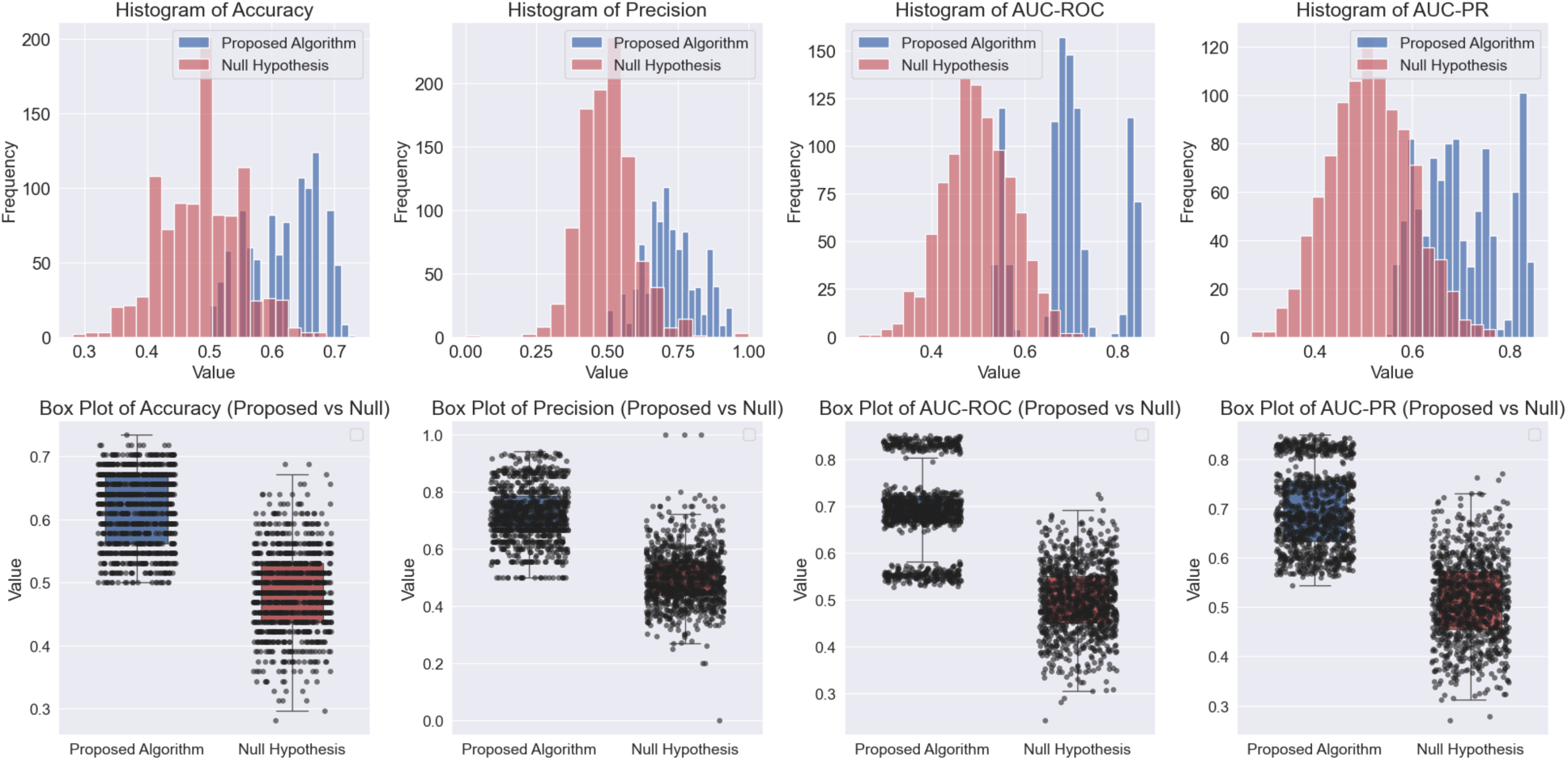
Comparison of the proposed algorithm and the null hypothesis for classification performance. The top row presents histograms of accuracy, precision, AUC-ROC, and AUC-PR, highlighting the distribution of results. The bottom row shows corresponding box plots, illustrating differences in median values and variability.

**Figure 9.**
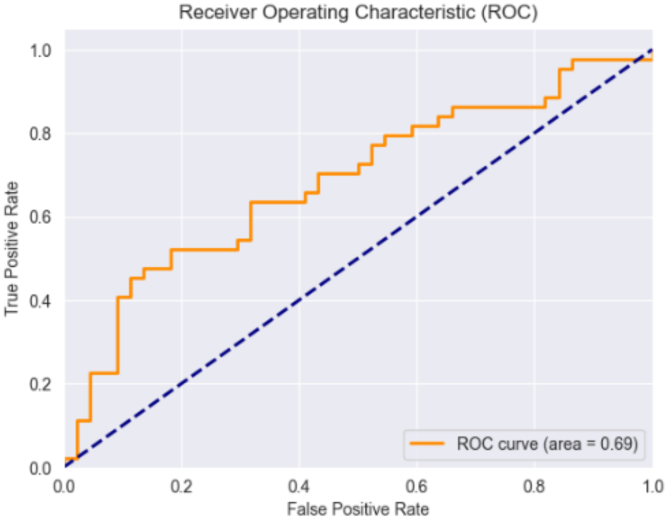
The receiver operating curve of the unseen data (Out-of-Sample ROC) The curve shows the trade-off between the TPR and FPR at various threshold settings. The dashed diagonal line represents the baseline.

## 4. Discussion

The results of this study demonstrate that our machine learning pipeline, which includes PCA and linear regression, can reliably and validly predict subjective craving intensity based on objective drug cue reactivity fMRI data. These results enhance our understanding of the neurobiological contributions to craving and underscore their clinical relevance for addiction and related disorders.

The comparable metrics for both in-sample and out-of-sample data underscore the generalizability of our model. While the effect sizes were in the moderate to large range, the statistical power was not consistently high, suggesting that additional data may be necessary for more robust validation. The positive trend in the slope of the lines connecting minimum and maximum craving points suggests that the model captures the variations in craving intensity effectively (Supplementary Figure 3). This reliable differentiation between extremes in craving levels supports the potential application of the model in understanding and predicting craving dynamics in substance use disorders.

By identifying the brain activity map, specifically the roles of the Parahippocampal Gyrus, Superior Temporal Gyrus, Medioventral Occipital Cortex, Amygdala (positive contribution), and Inferior Temporal Gyrus (negative contribution), we provide valuable insights into the brain regions that influence craving intensity. The Parahippocampal Gyrus and Amygdala play a well-established role in craving, as these regions are crucial for emotional processing, memory, and responses to cues—all central mechanisms in craving (Breiter et al., 1997; Koob & Volkow, 2010)). Their involvement aligns with findings in the literature, particularly in addiction studies. Superior Temporal Gyrus is involved in sensory integration and emotional regulation (Bigler et al., 2007; Radua et al., 2010), which can also contribute to craving responses. Medioventral Occipital Cortex is linked to visual processing, potentially related to visual cues triggering craving (Kanwisher & Yovel, 2006; Sabatinelli et al., 2009). The negative contribution of Inferior Temporal Gyrus (ITG) in craving could reflect inhibitory or regulatory processes, but this might require further investigation to align with existing studies. The ITG is primarily associated with visual object recognition and processing within the ventral visual stream (Haxby et al., 1996). Its involvement in craving may be linked to the processing of visual cues related to addictive substances. However, the specific role of the ITG in inhibitory control over craving is not well-established and needs further investigation.

The algorithm developed in (Koban et al., 2023) identified key brain regions associated with craving across multiple groups, including individuals with alcohol use disorder, cocaine use disorder, and cigarette. These regions included the ventromedial prefrontal and cingulate cortices, ventral striatum, temporal/parietal association areas, mediodorsal thalamus and cerebellum. These findings align with our results as both highlight temporal regions suggesting a shared emphasis on regions involved in memory and sensory processing. These functions are crucial for cue-induced cravings, as memory and sensory cues can trigger cravings. On the other hand, while they emphasize higher-order areas like the ventromedial prefrontal cortex, cingulate cortices, and ventral striatum, which are involved in decision-making, emotional regulation, and reward processing, our results highlight the amygdala, Parahippocampal Gyrus, and occipital cortex, indicating a stronger focus on emotional response (amygdala) and visual processing (medioventral occipital cortex) related to visual cue-induced craving.

The developed classifiers for discriminating between high and low craving levels and also distinguishing between neutral and drug-related cues based on fMRI data highlights their potential utility in clinical settings. These capabilities could inform the development of targeted interventions and personalized treatment plans for individuals with substance use disorders.

To enhance our findings, increasing access to larger datasets could significantly improve the generalizability of results, allowing for more nuanced conclusions. Incorporating additional covariates, including sex, weight, age, education level, handedness, and socioeconomic status, is essential to reduce potential confounding variables, thereby improving model fit and validity. Furthermore, employing a self-regulation strategy during data analysis could provide insights into how individual differences influence craving responses and modulation, ultimately yielding a more comprehensive understanding of the underlying mechanisms (Muraven & Baumeister, 2000).

Moreover, expanding our investigation to consider the effects of various other substances will allow for a comparative analysis that may uncover unique neurobiological signatures associated with different drugs. Lastly, exploring the impact of food and other sensory cues on craving could illuminate how environmental factors interact with individual neurobiology to shape craving intensity (Field & Cox, 2008). Collectively, these approaches not only promise to enhance our findings but also contribute to a more holistic understanding of craving in the context of addiction research.

## 5. Conclusion

We developed a machine learning pipeline that is able to predict the subjective craving based on objective fMRI image. The process involved multiple key steps: first, optimizing the model based on brain functional activity data, followed by identifying the most critical brain activity map. We then categorized the craving levels and cue types based on this pipeline. A vital part of the analysis was ensuring the generalization and statistical significance of the results across participants.

In terms of the identified activity map, several regions showed the most significant impact on craving intensity. The Parahippocampal Gyrus, Superior Temporal Gyrus, Medioventral Occipital Cortex, and Amygdala emerged as having the most positive contribution to craving. Conversely, the Inferior Temporal Gyrus showed a negative contribution. This comprehensive pipeline allows for the classification of brain activity patterns related to craving, advancing our understanding of the underlying neurobiological mechanisms. Overall, our study represents a significant step towards objective, neurobiologically-informed assessments of craving, with implications for improving addiction treatment and outcomes. For further validation, the model should be evaluated in varied populations and settings.

## Acknowledgements

We are grateful to Prof. Monica Luciana from the Department of Psychology for her support and valuable insights. We also acknowledge Rayus Kuplicki for providing the data essential to this research and for offering insightful contributions. Additionally, we appreciate the support from the Laureate Institute for Brain Research for sharing the database and providing insights into its data analytics.

## References

Addiction Cue-Reactivity Initiative (ACRI) Network, Sangchooli, A., Zare-Bidoky, M., Fathi Jouzdani, A., Schacht, J., Bjork, J. M., Claus, E. D., Prisciandaro, J. J., Wilson, S. J., Wüstenberg, T., Potvin, S., Ahmadi, P., Bach, P., Baldacchino, A., Beck, A., Brady, K. T., Brewer, J. A., Childress, A. R., Courtney, K. E., … Ekhtiari, H. (2024). Parameter Space and Potential for Biomarker Development in 25 Years of fMRI Drug Cue Reactivity: A Systematic Review. JAMA Psychiatry, 81(4), 414. 10.1001/jamapsychiatry.2023.5483

Anderson, T. W. (1958). An Introduction to Multivariate Statistical Analysis. Wiley.

Antons, S., Yip, S. W., Lacadie, C. M., Dadashkarimi, J., Scheinost, D., Brand, M., & Potenza, M. N. (2023). Connectome-based prediction of craving in gambling disorder and cocaine use disorder. Dialogues Clin. Neurosci., 25(1), 33–42.

Bigler, E. D., Mortensen, S., Neeley, E. S., Ozonoff, S., Krasny, L., Johnson, M., Lu, J., Provencal, S. L., McMahon, W., & Lainhart, J. E. (2007). Superior Temporal Gyrus, Language Function, and Autism. Developmental Neuropsychology, 31(2), 217–238. 10.1080/87565640701190841

Breiman, L. (2001). Random forests. Machine Learning, 45(1), 5–32. 10.1023/A:1010933404324

Chan, Y.-H., Chang, H.-M., Lu, M.-L., & Goh, K. K. (2024). Targeting cravings in substance addiction with transcranial direct current stimulation: Insights from a meta-analysis of sham-controlled trials. Psychiatry Research, 331, 115621. 10.1016/j.psychres.2023.115621

Chen, J., Qin, J., He, Q., & Zou, Z. (2020). A Meta-Analysis of Transcranial Direct Current Stimulation on Substance and Food Craving: What Effect Do Modulators Have? Frontiers in Psychiatry, 11, 598. 10.3389/fpsyt.2020.00598

Chen, R., Classen, J., Gerloff, C., Celnik, P., Wassermann, E. M., Hallett, M., & Cohen, L. G. (1997). Depression of motor cortex excitability by low-frequency transcranial magnetic stimulation. Neurology, 48(5), 1398–1403. 10.1212/WNL.48.5.1398

Chen, T., & Guestrin, C. (2016). XGBoost: A Scalable Tree Boosting System. Proceedings of the 22nd ACM SIGKDD International Conference on Knowledge Discovery and Data Mining, 785–794. 10.1145/2939672.2939785

Ekhtiari, H., Faghiri, A., Oghabian, M.-A., & Paulus, M. P. (2016). Functional neuroimaging for addiction medicine. In Progress in Brain Research (Vol. 224, pp. 129–153). Elsevier. 10.1016/bs.pbr.2015.10.001

Ekhtiari, H., Zare-Bidoky, M., Sangchooli, A., Janes, A. C., Kaufman, M. J., Oliver, J. A., Prisciandaro, J. J., Wüstenberg, T., Anton, R. F., Bach, P., Baldacchino, A., Beck, A., Bjork, J. M., Brewer, J., Childress, A. R., Claus, E. D., Courtney, K. E., Ebrahimi, M., Filbey, F. M., … Zilverstand, A. (2022). A methodological checklist for fMRI drug cue reactivity studies: Development and expert consensus. Nature Protocols, 17(3), 567–595. 10.1038/s41596-021-00649-4

Fan, L., Li, H., Zhuo, J., Zhang, Y., Wang, J., Chen, L., Yang, Z., Chu, C., Xie, S., Laird, A. R., Fox, P. T., Eickhoff, S. B., Yu, C., & Jiang, T. (2016). The Human Brainnetome Atlas: A New Brain Atlas Based on Connectional Architecture. Cerebral Cortex, 26(8), 3508– 3526. 10.1093/cercor/bhw157

Field, M., & Cox, W. (2008). Attentional bias in addictive behaviors: A review of its development, causes, and consequences. Drug and Alcohol Dependence, 97(1–2), 1–20. 10.1016/j.drugalcdep.2008.03.030

Garrison, K. A., Sinha, R., Potenza, M. N., Gao, S., Liang, Q., Lacadie, C., & Scheinost, D. (2023). Transdiagnostic Connectome-Based Prediction of Craving. American Journal of Psychiatry, 180(6), 445–453. 10.1176/appi.ajp.21121207

Hanlon, C. A., Dowdle, L. T., Austelle, C. W., DeVries, W., Mithoefer, O., Badran, B. W., & George, M. S. (2015). What goes up, can come down: Novel brain stimulation paradigms may attenuate craving and craving-related neural circuitry in substance dependent individuals. Brain Research, 1628, 199–209. 10.1016/j.brainres.2015.02.053

Haxby, J. V., Ungerleider, L. G., Horwitz, B., Maisog, J. M., Rapoport, S. I., & Grady, C. L. (1996). Face encoding and recognition in the human brain. Proceedings of the National Academy of Sciences, 93(2), 922–927. 10.1073/pnas.93.2.922

Hoerl, A. E., & Kennard, R. W. (1970). Ridge Regression: Biased Estimation for Nonorthogonal Problems. Technometrics, 12(1), 55–67. 10.1080/00401706.1970.10488634

Jansen, J. M., Van Wingen, G., Van Den Brink, W., & Goudriaan, A. E. (2015). Resting state connectivity in alcohol dependent patients and the effect of repetitive transcranial magnetic stimulation. European Neuropsychopharmacology, 25(12), 2230–2239. 10.1016/j.euroneuro.2015.09.019

Kanwisher, N., & Yovel, G. (2006). The fusiform face area: A cortical region specialized for the perception of faces. Philosophical Transactions of the Royal Society B: Biological Sciences, 361(1476), 2109–2128. 10.1098/rstb.2006.1934

Kilts, C. D., Gross, R. E., Ely, T. D., & Drexler, K. P. G. (2004). The Neural Correlates of Cue-Induced Craving in Cocaine-Dependent Women. American Journal of Psychiatry, 161(2), 233–241. 10.1176/appi.ajp.161.2.233

Koban, L., Wager, T. D., & Kober, H. (2023). A neuromarker for drug and food craving distinguishes drug users from non-users. Nat. Neurosci., 26(2), 316–325.

Kohavi, R. (1995). A study of cross-validation and bootstrap for accuracy estimation and model selection. In Proceedings of the 14th International Joint Conference on Artificial Intelligence (pp. 1137–1143).

Koob, G. F., & Volkow, N. D. (2010). Neurocircuitry of Addiction. Neuropsychopharmacology, 35(1), 217–238. 10.1038/npp.2009.110

Kuhn, J., Möller, M., Treppmann, J. F., Bartsch, C., Lenartz, D., Gruendler, T. O. J., Maarouf, M., Brosig, A., Barnikol, U. B., Klosterkötter, J., & Sturm, V. (2014). Deep brain stimulation of the nucleus accumbens and its usefulness in severe opioid addiction. Molecular Psychiatry, 19(2), 145–146. 10.1038/mp.2012.196

Luigjes, J., Van Den Brink, W., Feenstra, M., Van Den Munckhof, P., Schuurman, P. R., Schippers, R., Mazaheri, A., De Vries, T. J., & Denys, D. (2012). Deep brain stimulation in addiction: A review of potential brain targets. Molecular Psychiatry, 17(6), 572–583. 10.1038/mp.2011.114

Muraven, M., & Baumeister, R. F. (2000). Self-regulation and depletion of limited resources: Does self-control resemble a muscle? Psychological Bulletin, 126(2), 247–259. 10.1037/0033-2909.126.2.247

Naqvi, N. H., & Bechara, A. (2010). The insula and drug addiction: An interoceptive view of pleasure, urges, and decision-making. Brain Structure and Function, 214(5–6), 435–450. 10.1007/s00429-010-0268-7

Pedregosa, F., Varoquaux, G., & Gramfort, A. (2011). Scikit-learn: Machine Learning in Python. JMLR.Org.

Pettorruso, M., Di Giannantonio, M., De Risio, L., Martinotti, G., & Koob, G. F. (2020). A Light in the Darkness: Repetitive Transcranial Magnetic Stimulation (rTMS) to Treat the Hedonic Dysregulation of Addiction. Journal of Addiction Medicine, 14(4), 272–274. 10.1097/ADM.0000000000000575

Pollard, A. A., Hauson, A. O., Lackey, N. S., Zhang, E., Khayat, S., Carson, B., Fortea, L., Radua, J., & Grant, I. (2023). Functional neuroanatomy of craving in heroin use disorder: Voxel-based meta-analysis of functional magnetic resonance imaging (fMRI) drug cue reactivity studies. The American Journal of Drug and Alcohol Abuse, 49(4), 418–430. 10.1080/00952990.2023.2172423

Radua, J., Phillips, M. L., Russell, T., Lawrence, N., Marshall, N., Kalidindi, S., El-Hage, W., McDonald, C., Giampietro, V., Brammer, M. J., David, A. S., & Surguladze, S. A. (2010). Neural response to specific components of fearful faces in healthy and schizophrenic adults. NeuroImage, 49(1), 939–946. 10.1016/j.neuroimage.2009.08.030

Sabatinelli, D., Lang, P. J., Bradley, M. M., Costa, V. D., & Keil, A. (2009). The Timing of Emotional Discrimination in Human Amygdala and Ventral Visual Cortex. The Journal of Neuroscience, 29(47), 14864–14868. 10.1523/JNEUROSCI.3278-09.2009

Shen, Z., Huang, P., Wang, C., Qian, W., Yang, Y., & Zhang, M. (2017). Increased network centrality as markers of relapse risk in nicotine-dependent individuals treated with varenicline. Prog. Neuropsychopharmacol. Biol. Psychiatry, 75, 142–147.

Sinha, R. (2013). The clinical neurobiology of drug craving. Current Opinion in Neurobiology, 23(4), 649–654. 10.1016/j.conb.2013.05.001

Tibshirani, R. (1996). Regression Shrinkage and Selection Via the Lasso. Journal of the Royal Statistical Society Series B: Statistical Methodology, 58(1), 267–288. 10.1111/j.2517-6161.1996.tb02080.x

Tiffany, S. T., & Wray, J. M. (2012). The clinical significance of drug craving. Annals of the New York Academy of Sciences, 1248(1), 1–17. 10.1111/j.1749-6632.2011.06298.x

Volkow, N. D., Wang, G.-J., Fowler, J. S., Logan, J., Gatley, S. J., Hitzemann, R., Chen, A. D., Dewey, S. L., & Pappas, N. (1997). Decreased striatal dopaminergic responsiveness in detoxified cocaine-dependent subjects. Nature, 386(6627), 830–833. 10.1038/386830a0

Weiss, R. D., Griffin, M. L., & Hufford, C. (1995). Craving in Hospitalized Cocaine Abusers as a Predictor of Outcome. The American Journal of Drug and Alcohol Abuse, 21(3), 289–301. 10.3109/00952999509002698

Zilverstand, A., Huang, A. S., Alia-Klein, N., & Goldstein, R. Z. (2018). Neuroimaging Impaired Response Inhibition and Salience Attribution in Human Drug Addiction: A Systematic Review. Neuron, 98(5), 886–903. 10.1016/j.neuron.2018.03.048

Zou, H., & Hastie, T. (2005). Regularization and Variable Selection Via the Elastic Net. Journal of the Royal Statistical Society Series B: Statistical Methodology, 67(2), 301–320. 10.1111/j.1467-9868.2005.00503.x

