## Supplementary Material for "Machine Learning-Driven fMRI Analysis for Objective Craving Prediction"

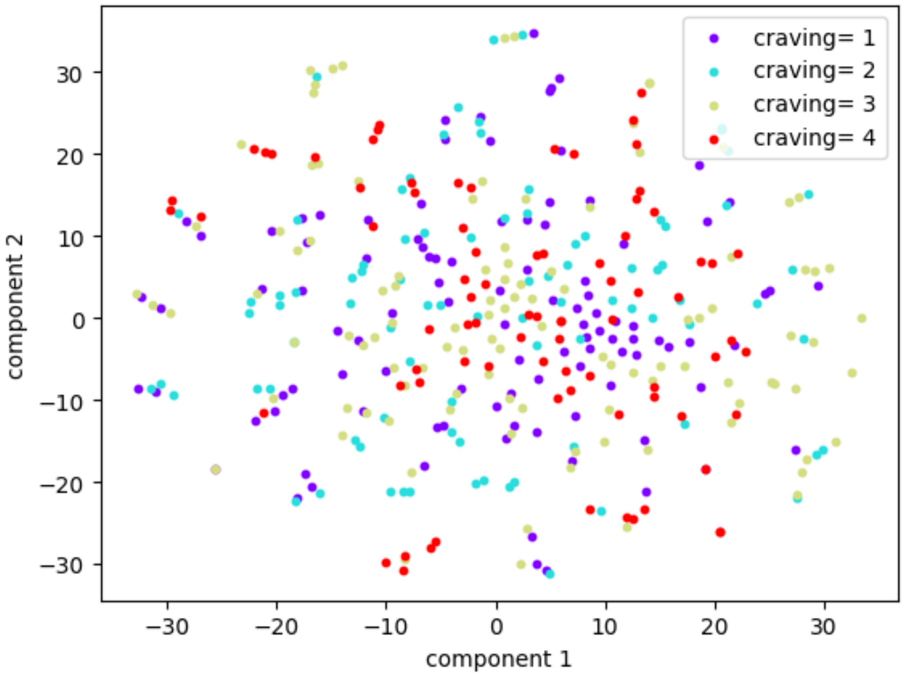


**Supplementary Figure 1. Visualization of the First Two Components of t-SNE (t-Distributed Stochastic Neighbor Embedding) applied to fMRI data.** This 2D representation shows the reduced-dimensional structure of fMRI beta coefficients derived from cue-reactivity tasks. Each point represents a sample, with different colors indicating distinct craving levels. The t-SNE algorithm preserves local relationships, allowing for a qualitative assessment of how neural responses relate to craving intensity.

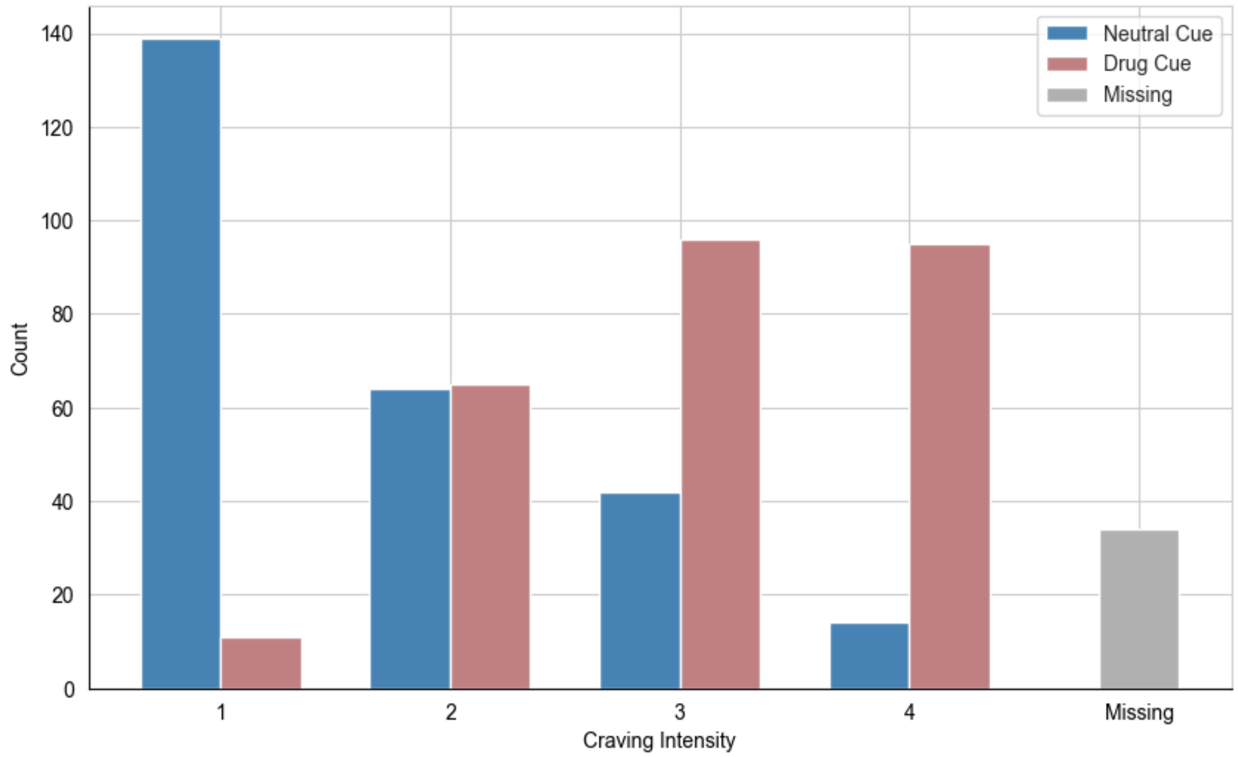


**Supplementary  Figure 2. Distribution of self-reported craving intensity for neutral and drug cues, rated on a scale from 1 to 4.** The overall mean craving intensity is 2.37 ± 1.10 (Std Dev), reflecting moderate variability in responses. The median value of 2.0 indicates that half of the responses fall below this level, emphasizing a tendency toward lower craving levels. Neutral cues elicited lower craving ratings (1.76 ± 0.91), while drug cues resulted in higher reported cravings (3.02 ± 0.86), demonstrating a clear distinction between conditions. There are 42 missing values, accounting for 7.5% of the dataset.

| Feature Selection | Regression Model | Parameters | RMSE +/- Std |
|---|---|---|---|
| PCA | Linear Regression | pca__n_components': 100 | 0.972 +/- 0.016 |
| PCA | Lasso | pca__n_components': 100, reg_alpha': 0.9 | 0.972 +/- 0.011 |
| PCA | Elastic Net | pca__n_components': 100, reg_alpha': 0.1, reg_l1_ratio': 0.9 | 0.971 +/- 0.007 |
| PCA | Random Forest | pca__n_components': 180, reg_max_depth': None, reg_min_samples_split': 10, reg_n_estimators': 50 | 1.009 +/- 0.067 |
| PCA | XGBoost | pca__n_components': 180, reg_learning_rate': 0.01, reg_max_depth': 3, reg_n_estimators': 50 | 0.996 +/- 0.009 |
| ANOVA (K Best) | Linear Regression | select_k_best__k': 2 | 1.057 +/- 0.077 |
| ANOVA (K Best) | Lasso | reg_alpha': 1, select_k_best__k': 10 | 0.998 +/- 0.003 |
| ANOVA (K Best) | Elastic Net | reg_alpha': 0.9, reg_l1_ratio': 0.1, select_k_best__k': 2 | 0.998 +/- 0.023 |
| ANOVA (K Best) | Random Forest | reg_max_depth': None, reg_min_samples_split': 10, reg_n_estimators': 100, select_k_best__k': 2 | 1.126 +/- 0.070 |
| ANOVA (K Best) | XGBoost | reg_learning_rate': 0.01, reg_max_depth': 3, reg_n_estimators': 50, select_k_best__k': 2 | 1.006 +/- 0.032 |

**Supplementary Table 1. Hyperparameter Tuning Results.** The optimization results aimed at minimizing RMSE were obtained by evaluating different combinations of feature selection methods, regression models and their respected hyperparameters.

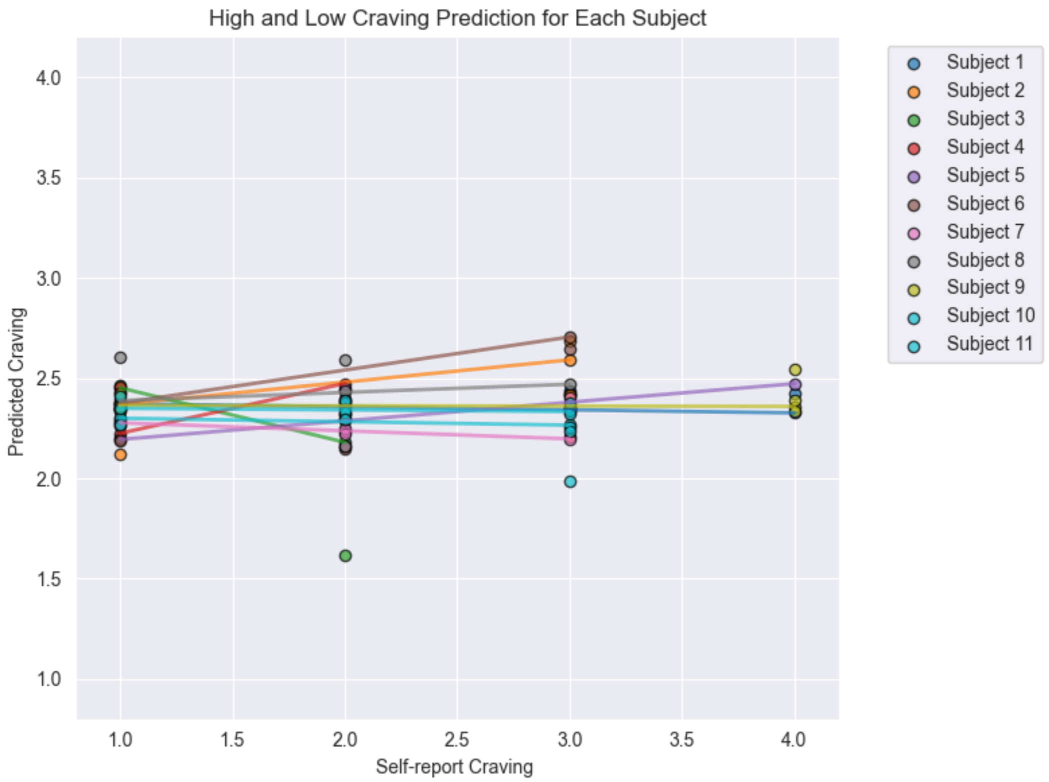


**Supplementary Figure 3. Out-of-sample results for PCA (n=100) and linear regression.** The lines connect the points representing the lowest craving on the x-axis and its predicted value on the y-axis to the points representing the highest craving on the x-axis and its predicted value on the y-axis for each subject in the out-of-sample set.
